# Simultaneous loss of CAMK2A and CAMK2B reveals endogenous in vivo substrates

**DOI:** 10.1101/2024.11.17.624016

**Authors:** Pomme M.F. Rigter, Karel Bezstarosti, Oguz Can Koc, Tyler L. Perfitt, Jeroen A.A. Demmers, Roger J. Colbran, Margaret Stratton, Ype Elgersma, Geeske M. van Woerden

## Abstract

Ca^2+^/calmodulin-dependent protein kinase 2 (CAMK2) plays a critical role in calcium signaling. Recent gene knockout studies show that CAMK2A and CAMK2B can have distinct roles yet also partially compensate for each other in yet unknown brain functions. In order to provide insight into potential novel CAMK2 functions, we performed parallel phosphoproteomic analyses on non-stimulated cortex tissue from inducible *Camk2a* and *Camk2b* double knockout (*Camk2a^f/f^;Camk2b^f/f^;CAG-Cre^ESR^*) mice and from wild type mice. A total of 5622 phosphorylated peptides derived from 2080 proteins were identified. Phosphorylation at serine/threonine residues in 130 proteins were downregulated in the double knockout mice, including residues in 113 proteins that have not previously been identified as potential CAMK2 substrates. Comparison of amino acid sequences surrounding the downregulated phosphorylation residues provided new insights into the CAMK2-substrate consensus sequences *in vivo*. This dataset provides an important resource for future studies examining novel roles for CAMK2 in the brain.

## Introduction

One of the most remarkable functional properties of the brain is synaptic plasticity, the process by which novel experiences elicit changes in neuronal network formation and activation to form new memories. The Ca^2+^ second messenger is a critical mediator of these processes, in large part by regulating Ca^2+^/calmodulin-dependent protein kinase 2 (CAMK2). CAMK2 consists of a family of four proteins CAMK2A (alpha), CAMK2B (beta), CAMKG (gamma) and CAMK2D (delta) that share high homology and are encoded by individual genes. CAMK2A and CAMK2B have been most intensively studied and comprise 2% of total protein in the hippocampus, 1.3% in the cortex (CAMK2A) (Erondu & Kennedy, 1985), and up to 7.4% (CAMK2A) and 1.3% (CAMK2B) of total protein in the postsynaptic density (PSD) (Cheng et al., 2006). They form and function in homomeric or heteromeric holoenzymes consisting of 12-14 subunits with an average 3:1 (CAMK2A:CAMK2B) ratio in the forebrain (Bennett et al., 1983; Bhattacharyya et al., 2016; Brocke et al., 1999; Myers et al., 2017; Vallano, 1989). It is well established that CAMK2 translates transient increases in postsynaptic calcium levels following neuronal activation into long-lasting effects via autophosphorylation at position Thr286 in the alpha isozyme and position Thr287 in other isozymes (Bayer & Schulman, 2019; Hell, 2014; Hudmon & Schulman, 2002).

Even though CAMK2A and CAMK2B share a high homology (84.1% amino acid sequence identity in mouse) and function together in heteromeric holoenzymes, they have unique structural domains and protein functions. The use of *Camk2a* and *Camk2b* knockout and point mutant mouse models altering enzymatic activity has shown that for synaptic plasticity and learning in mouse hippocampus the enzymatic function of CAMK2A is critical, whereas CAMK2B has a more structural role (Achterberg et al., 2014; Borgesius et al., 2011; Elgersma et al., 2002; Giese et al., 1998; Silva, Paylor, et al., 1992; Silva, Stevens, et al., 1992). However, we recently showed that simultaneous deletion or silencing of the enzymatic activity of CAMK2A and CAMK2B is lethal (Kool et al., 2019). This suggests that CAMK2A and CAMK2B can compensate for each other in controlling essential brain functions that remain to be uncovered. More extensive knowledge about this redundancy, for example by looking at the full list of substrates of CAMK2, could provide new understanding on the full functional spectrum of CAMK2 in the brain.

Synapsin I was the first CAMK2 substrate identified (Bennett et al., 1983), and many other substrates have since been identified. A majority of these substrates were identified using purified brain CAMK2 containing a mixture of different isozymes, or using purified CAMK2A (Fink & Meyer, 2002; Shonesy et al., 2014). The high homology in sequence identity in the catalytic domains of all the CAMK2 proteins suggests that the other variants have a very similar substrate specificity. In addition, since CAMK2A and CAMK2B can compensate for each other, it is likely that additional common substrates remain to be identified. Moreover, the specific site(s) phosphorylated in many CAMK2 substrates remain undetermined.

An extensive list of established and putative interaction partners, termed CAMK2 associated proteins (CAMKAPs), have been identified using proteomics (Baucum et al., 2015). Additionally, databases have been generated on the brain phosphoproteome, most likely describing CAMK2-dependent phosphorylation sites (Goswami et al., 2012; Huttlin et al., 2010; Jaffe et al., 2004; Li et al., 2016; Lundby et al., 2012; Trinidad et al., 2012). Only recently the first phosphoproteomics study on NMDA receptor-dependent activity in the striatum upon *ex vivo* stimulation was published, revealing an extensive list of substrates consistent with the consensus motif for CAMK2-dependent phosphorylation (Funahashi et al., 2024). However, the impact of genetic deletion of CAMK2 on the neuronal phosphoproteome *in vivo* has not yet been investigated. In addition, as CAMK2A and CAMK2B can partially compensate for each other (Kool et al., 2019), it is likely that still unknown CAMK2 substrates exist. In order to gain further insight in the understanding how phosphorylation changes in the absence of enzymatic activity of CAMK2A and CAMK2B, we compared the phosphoproteomes from unstimulated cortical tissue of wild type and *Camk2a^f/f^;Camk2b^f/f^;CAG-Cre^ESR^*mice by performing a TMT labeling based phosphoproteomic analyses. We characterized CAMK2 autophosphorylation sites and identified novel potential substrates of CAMK2, providing new insights into the consensus CAMK2 phosphorylation motif. Together, our results provide a valuable resource for future research on the role of CAMK2 in the brain.

## Materials and methods

### Animals

In this study we tested both sexes of previously described *Camk2a^f/f^;Camk2b^f/f^;CAG-Cre^ESR^* (Kool et al., 2019), *Camk2a^-/-^* (Achterberg et al., 2014; Elgersma et al., 2002) and *Camk2b^-/-^*mouse lines (Borgesius et al., 2011; Kool et al., 2016) in C57BL/6J background backcrossed >16 times. Mice were group housed in IVC cages (Sealsafe 1145T, Tecniplast) and kept on light/dark 12/12h cycle at 21±2 °C with food and water available *ad libitum*. All experiments were done during the light phase and experimenters were blind to genotypes. Mice were genotyped between 7-10 days of age and re-genotyped after the mice were sacrificed. Genotyping records were obtained and kept by a technician not involved in the design, execution or analysis of the experiments.

All animal experiments were conducted in accordance with the European Commission Council Directive 2010/63/EU (CCD project license AVD101002017893), and all described experiments and protocols were subjected to ethical review and approved by an independent review board of the Erasmus MC.

### Tamoxifen

For Cre-mediated deletions, *Camk2a^f/f^;Camk2b^f/f^* and *Camk2a^f/f^;Camk2b^f/f^;CAG-Cre^ESR^* mice received intraperitoneal injections with 0.1 mg tamoxifen (Sigma-Aldrich) per gram bodyweight for four consecutive days. Tamoxifen was prepared fresh by dissolving it in sunflower oil at a concentration of 20 mg/ml. To keep the dose constant throughout injection days we injected mice in the afternoon, 24±1 hours after the previous injection.

### Mass Spectrometry

Samples were prepared from adult *Camk2a^f/f^;Camk2b^f/f^* and *Camk2a^f/f^;Camk2b^f/f^;CAG-Cre^ESR^*mice (8–10 weeks of age, n = 3) that were injected with tamoxifen for four consecutive days and were sacrificed 21 days after the first injection. Cortical tissue was isolated and lysed in 1 ml 50 mM Tris/HCl pH 8.2, 0.5 % sodium deoxycholate (SDC) and MS-SAFE™ protease and phosphatase inhibitor using a Bioruptor ultrasonicator (Diagenode). Protein concentrations were measured using the BCA assay (Thermo Scientific). Proteins were reduced with 5 mM DTT and cysteine residues were alkylated with 10 mM iodoacetamide. Protein was extracted by acetone precipitation at -20 °C overnight. Samples were centrifuged at 8,000 g for 10 min at 4 °C. The acetone was removed and the pellet allowed to dry. The protein pellet (∼1 mg protein) was dissolved in 1 ml 50 mM Tris/HCl pH 8.2, 0.5 % SDC and proteins were digested with LysC (1:200 enzyme:protein ratio) for 4 h at 37 °C. Next, trypsin was added (1:100 enzyme:protein ratio) and the digestion proceeded overnight at 30 °C. Digests were acidified with 50 μl 10 % formic acid (FA) and centrifuged at 8,000 g for 10 min at 4 °C to remove the precipitated SDC. The supernatant was transferred to a new centrifuge tube. The digests were purified with C18 solid phase extraction (Sep-Pak, Waters), lyophilized and stored at -20 °C.

Phosphopeptide enrichment was carried out according to Kettenbach et al. (Kettenbach et al., 2011) with some modifications. 1 mg of lyophilized peptide digest was dissolved in 1 ml 50 % acetonitrile (AcN), 2 M lactic acid with 6 mg TiO2 beads (GL Sciences) and incubated on a rotator at room temperature for 2 h. Beads were washed twice with 2 M lactic acid / 50 % AcN and once with 4% FA in 50 % AcN. Phosphopeptides were eluted twice with 150 μL of 50 mM K_2_HPO_4_, 1% pyrrolidine, acidified with 90 μL of 10 % FA and stored at -20 °C. Extracted proteolytic peptides were labeled with TMT 10-plex labeling reagents (Thermo Scientific) allowing for peptide quantitation. Peptides were mixed at the 16-plex level and further fractionated into six fractions by HILIC chromatography. Fractions were collected and analyzed by nanoflow LC-MS/MS. nLC-MS/MS was performed on EASY-nLC 1000 coupled to an Orbitrap Fusion Tribrid mass spectrometer (ThermoFisher Scientific) operating in positive mode and equipped with a nanospray source. Peptides were separated on a ReproSil C18 reversed phase column (Dr Maisch GmbH; column dimensions 15 cm × 50 µm, packed in-house) using a linear gradient from 0 to 80% B (A = 0.1 % formic acid; B = 80% (v/v) acetonitrile, 0.1 % formic acid) in 70 min and at a constant flow rate of 200 nl/min using a splitter. The column eluent was directly sprayed into the ESI source of the mass spectrometer. Mass spectra were acquired in continuum mode; fragmentation of the peptides was performed in data-dependent mode using the multinotch SPS MS3 reporter ion-based quantification method.

Data were analyzed with Proteome Discoverer 2.1. Peak lists were created from raw data files using the Mascot Distiller software (version 2.3; MatrixScience). The Mascot search algorithm (version 2.3.2, MatrixScience) was used for searching against the Uniprot database (taxonomy: *Mus musculus*, version December 2015). The peptide tolerance was typically set to 10 ppm and the fragment ion tolerance was set to 0.8 Da. The reporter ion tolerance was set to 0.003 Da. A maximum number of 2 missed cleavages by trypsin were allowed and carbamidomethylated cysteine was set as fixed modifications, while oxidized methionine and TMT on lysine and N-terminus were set as variable modifications. Typical contaminants were omitted from the output tables. Protein ratios were calculated from the scaled normalized abundances of the reporter ions over the six quantitation channels. Only phosphopeptides with confidence scores ‘high’ and ‘medium’ were used for further calculations. Proteins corresponding to phosphospeptides that were >1.5 fold significantly downregulated in *Camk2a^f/f^;Camk2b^f/f^;CAG-Cre^ESR^*compared to *Camk2a^f/f^;Camk2b^f/f^* samples were used as the input for GO enrichment analysis using open resource Metascape (Zhou et al., 2019; https://metascape.org/). Settings were adapted to *Mus musculus*, and the background list to all phosphoproteins identified in our phosphoproteome screen data. Enrichment analysis was done on GO terms for biological processes, cellular components and molecular functions with adapted P-values using Benjamin-Hochberg procedure and hierarchical clustering using Kappa scores. Phosphosites were mapped on CAMK2A (Uniprot accession P11798) and CAMK2B (Uniprot accession Q5SVJ0). For the motif analysis, phosphosites were set at position 0 and sequences were manually supplemented to 10 residues on both sides, amino acid abundance was visualized using WebLogo (http://weblogo.berkeley.edu/). To determine if our list of potential CAMK2 substrates were novel, the phosphospeptides that were >1.5 fold downregulated in *Camk2a^f/f^;Camk2b^f/f^;CAG-Cre^ESR^* samples were compared to the list of CAMK2A or CAMK2B substrates in all species on the PhosphoSitePlus^®^ PTM database (Hornbeck et al., 2015), which was downloaded on the 27^th^ of October 2024.

### Statistics

Statistics were performed in R, Microsoft Excel or Metascape, with α set at 0.05. Because our goal here was to execute an exploratory analysis, in which we aimed to generate hypotheses rather than to confirm them, we decided to not correct for the p-values of phoshopeptides downregulated >1.5 in *Camk2a^f/f^;Camk2b^f/f^;CAG-Cre^ESR^* mice. Subsequent analyses have been corrected. All values represent mean ± SEM. *p < 0.05, **p < 0.01, ***p < 0.001.

### CAMK2 kinase domain purification

Kinase domain of CAMK2 alpha (residues 7-274 from Uniprot: Q9UQM7) was purified as previously described (Ozden et al., 2022). CAMK2 was co-expressed with lambda phosphatase in *E. coli* BL21 (DE3) strain. Expression was induced by 1 mM IPTG and cells were grown at 18°C overnight. Following centrifugation, pellets were resuspended in Buffer A (25 mM Tris-HCl pH 8.5, 50 mM KCl, 40 mM Imidazole, 10% glycerol) supplemented with protease inhibitors (500 μM benzamidine, 200 μM AEBSF, 5 μM leupeptin, 1 μg/ml pepstatin, 100 μg/ml trypsin inhibitor), DNase (1 μg/ml), and 50 mM MgCl_2_. Cells were then lysed using a cell disruptor (Avestin, C50), and the lysates were centrifuged at 38,000 g at 4°C for 1 hour. Subsequently, lysates were loaded onto HisTrap FF columns (cytiva), washed with Buffer A, and eluted with Buffer B (25 mM Tris-HCl pH 8.5, 100 mM KCl, 520 mM Imidazole, 10% glycerol). Afterwards, the excess imidazole was removed from the sample using HiPrep 26/10 desalting column (cytiva) in Buffer C (25 mM Tris-HCl pH 8.5, 40 mM KCl, 40 mM Imidazole, 2 mM TCEP, 10% glycerol). 6xHis-SMT3 tag was cleaved from the protein by Ulp1 protease at 4°C, overnight. Ulp1 and cleaved 6xHis-SMT3 tags were removed from the sample using HisTrap FF columns and the remaining sample was further purified by anion exchange using HiTrap Q FF columns (cytiva). Protein was eluted from the column by a KCl gradient. And the fractions containing CAMK2 alpha kinase domain were concentrated using Amicon ultra centrifugal filter (Millipore) before injecting them into HiLoad 16/600 Superdex 75 pg gel filtration column. Column was run with Buffer GF (25 mM Tris pH 8.0, 150 mM KCl, 1 mM TCEP, 10% glycerol). Final protein product was concentrated, flash frozen, and kept at -80°C until use.

### In vitro kinase assay

Kinase activity assays were adapted from previous publications (Barker et al., 1995; Chao et al., 2010). The assay was conducted in 115 mM Tris, 150 mM KCl, 10 mM MgCl2, 2 mM ATP, 1 mM Phosphoenolpyruvate (Alfa Aesar), 0.2 mM NADH (Sigma), 10 units/mL Pyruvate kinase (Sigma), 30 units/mL Lactate dehydrogenase (Millipore Sigma), and varying concentrations of peptide substrates (Genscript). The final CAMK2A kinase domain concentration was 5 nM. The reactions were started by the addition of kinase to the reaction mix and the NADH fluorescence was measured using a Synergy H1 microplate reader (Biotek) at 450 nm at 30°C for 10 min with 20 sec intervals. NADH fluorescence was converted to concentration by a standard curve (R^2^=0.9891). The rate was obtained by calculating the maximum observed slope of each reaction. Data were fitted using the Michaelis-Menten equation in GraphPad Prism version 10.1.2.

### Radiolabeled phosphorylation assay with GST fusion protein

GST fusion proteins containing residues 1468-1740 of Shank3b (rat: UniProtKB: Q9JLU4 Isoform 2) (WT and with a Ser1511 to Ala mutation) were expressed in E. coli (BL21(DE3) and purified using glutathione-agarose, as described previously (Perfitt, Stauffer, et al., 2020). GST-Sk3(1468-1740) or GST alone were incubated at 30°C for 2 min with purified CAMK2A (Robison et al., 2005) (10 nM) in 50 mM HEPES, pH 7.5, 10 mM magnesium acetate, 2 mM CaCl_2_, 2 μM calmodulin, 1 μM dithiothreitol, 400 μM [γ-^32^P] ATP (700-1,000 c.p.m./pmol). Reactions were quenched by adding SDS, heated (70°C, 10 min) and then resolved by SDS-PAGE. Gels were stained with Coomassie (InstantBlue, VWR) and then exposed to X-ray film (Phenix). Stained gels and X-ray films were digitally scanned and bands were quantified using ImageJ. Phosphorylated protein signals on the X-ray films were normalized to the total protein signals from the corresponding gels. The normalized S1511A mutant protein phosphorylation was expressed as a percentage of normalized WT protein phosphorylation in each experiment.

## Results

### Absence of Camk2a and Camk2b alters the phosphoproteome

To provide a global, unbiased assessment of the pathways that involve CAMK2A and CAM2B, we made use of our previously generated inducible double knockout *Camk2a^f/f^;Camk2b^f/f^;CAG-Cre^ESR^*mice in C57BL/6J background (Kool et al., 2019). We started gene deletion by injecting tamoxifen (0.1 mg/g) for four consecutive days in adult mice (8-10 weeks) and sacrificed them 21 days later under basal conditions (**Figure 1A**). At this timepoint, we have previously shown that CAMK2A and CAMK2B protein levels reduced to <10% on western blot (Kool et al., 2019). We isolated cortical tissue and performed TMT multiplex tandem mass spectrometry on peptides of whole lysate. Proteomics and phosphoproteomics were performed simultaneously on the lysates; data from the proteomics analysis has been published previously (Kool *et al.,* 2019).

**Figure 1:**
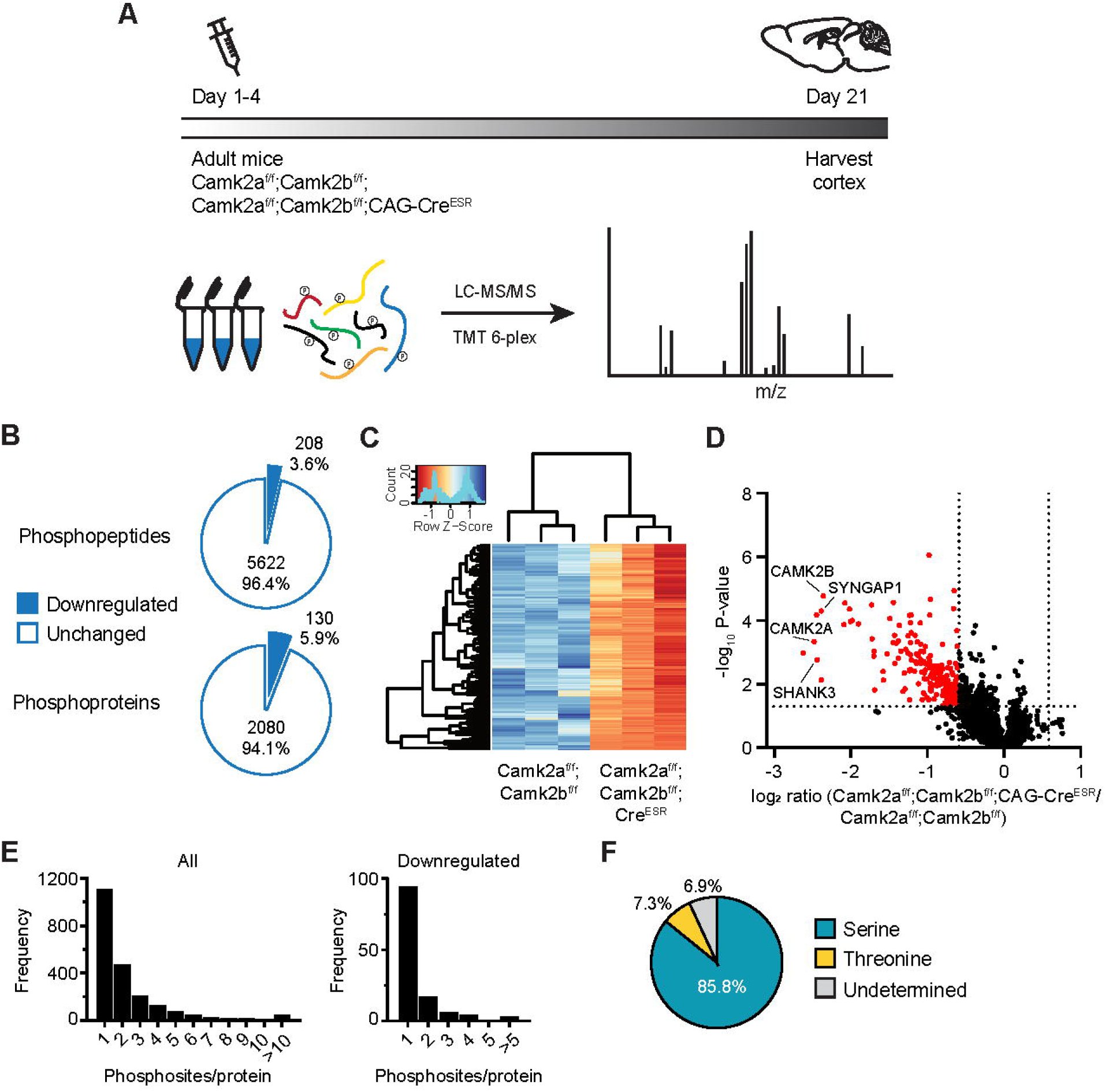
Phosphoproteomic dataset in *Camk2a^f/f^;Camk2b^f/f^;Cag-Cre^ESR^* mice. **(A)** To induce gene deletion, adult mutant *Camk2a^f/f^;Camk2b^f/f^;Cag-Cre^ESR^*and control *Camk2a^f/f^;Camk2b^f/f^* mice were injected with tamoxifen for four consecutive days and their brains were harvested 21 days later (n = 3). Experimental workflow for MS on cortical tissue. **(B)** Pie chart showing the amount and percentages of phosphorylated peptides (total 5830) and corresponding proteins (total 2210) that were >1.5 fold downregulated in *Camk2a^f/f^;Camk2b^f/f^;Cag-Cre^ESR^*mice. **(C)** Clustered heatmap of the downregulated phosphoproteins. **(D)** Volcano plot of all phosphopeptides, dashed lines indicate ratio difference at 1.5 and p-value at 0.05. Phosphopeptides significantly >1.5 downregulated are indicated in red. **(E)** Frequency distribution of number of phosphosites identified on the same protein, histogram of the complete dataset (left) and of the downregulated phosphopeptides in *Camk2a^f/f^;Camk2b^f/f^;Cag-Cre^ESR^* mice (right). **(F)** Phosphorylated amino acid distribution of CAMK2-dependent phosphosites.

From both *Camk2a^f/f^;Camk2b^f/f^;CAG-Cre^ESR^*and control *Camk2a^f/f^;Camk2b^f/f^* cortical samples, we enriched for phosphorylated peptides with titanium dioxide and identified 5830 phosphopeptides that corresponded to 2210 proteins (**Table S1**). Of these phosphopeptides, none were upregulated, whereas 208 phosphopeptides from a total of 130 proteins were significantly downregulated >1.5 fold in the *Camk2a^f/f^;Camk2b^f/f^;CAG-Cre^ESR^*samples, compared to *Camk2a^f/f^;Camk2b^f/f^* control (**Figure 1B-D** and **Table S2**), indicating multiple phosphosites on some of the substrates (**Figure 1E**). Proteins with multiple phosphosites include SYNGAP1 and the DLGAP family, which are known substrates of CAMK2. CAMK2 is a serine/threonine kinase, and previous results using peptides have shown that there is a small preference for serines over threonines (Johnson et al., 2023), though others have shown that replacing the serine with a threonine had little effect on CAMK2-dependent phosphorylation (Stokoe et al., 1993). In contrast, serine was the predominant phosphorylated residue that was suppressed in samples from *Camk2a^f/f^;Camk2b^f/f^;CAG-Cre^ESR^*mice (**Figure 1F**). The difference with previous published studies might result from differences in the sequences surrounding the target serines; for example, the amino acid immediately following the target residue can significantly affect phosphorylation (White et al., 1998).

Importantly, we found that the phosphorylation of several proteins/sites previously identified as CAMK2 substrates were significantly decreased in *Camk2a^f/f^;Camk2b^f/f^;CAG-Cre^ESR^*mice, including GRIN2A and GRIN2B, Calcineurin-subunit PPP3CB, SYN3, and LRRC7 (also known as Densin-180). The substrate showing the strongest reduction in phosphorylation in the *Camk2a^f/f^;Camk2b^f/f^;CAG-Cre^ESR^*mice, aside from CAMK2A and CAMK2B, was SHANK3 at Ser1510 (**Figure 1D**), indicating for the first time that this site is CAMK2-dependent in vivo (Dosemeci & Jaffe, 2010; Li et al., 2016; Wang et al., 2019). Among the new substrates detected in our experiment are additional key proteins involved in synaptic transmission and plasticity, such as IQSEC2, RPL6, ANK3, STX1B and KCNQ5, of which the first two are known CAMKAPs but here now also identified as phosphorylation substrates.

### Phosphorylation residues on CAMK2

As previously reported (Kool *et al.,* 2019), the proteomics analysis confirmed the successful deletion of CAMK2A and CAMK2B in in the *Camk2a^f/f^;Camk2b^f/f^;CAG-Cre^ESR^* mice, but the expression of very few other proteins was affected (**Fig S1**). Three weeks after onset of gene deletion, CAMK2A and CAMK2B protein levels in *Camk2a^f/f^;Camk2b^f/f^;CAG-Cre^ESR^* mice were very low compared to control *Camk2a^f/f^;Camk2b^f/f^* mice, but the levels of CAMK2G were unaffected and of CAMK2D only mildly but significantly affected (p = 0.0216, Unpaire 2-sided t-test) (**Figure 2A**). Being a substrate for itself, we analyzed which residues on CAMK2 were phosphorylated in our dataset. Because the isozymes are highly homologous to each other, some the sequences of some phosphopeptides derived from multiple CAMK2 proteins are identical (indicated in grey in **Figure 2B**). For example, the phosphopeptide QETVDC (Thr286) could be derived from CAMK2A or CAMK2D, although the levels of CAMK2D in these samples are substantially lower than those of CAMK2A. These data further confirm the *in vivo* identity of several previously identified CAMK2A and CAMK2B phosphorylation sites (Baucum et al., 2015; Huttlin et al., 2010; Trinidad et al., 2012). These residues are mainly located in the regulatory and linker domain (**Figure 2E**). Two residues (Ser279 and Thr330) were exclusively found to be phosphorylated in CAMK2A and three residues (T366, S367 and S371) in the unique linker domain of CAMK2B, of which S371 is involved in F-actin binding (Kim et al., 2015). As expected, we detected much lower levels of the phosphorylated peptides derived from CAMK2A (**Figure 2B**) and CAMK2B (**Figure 2C**) in samples from *Camk2a^f/f^;Camk2b^f/f^;CAG-Cre^ESR^*mice. However, while the reduced levels of some phosphorylated peptides were comparable to the loss of the corresponding protein (e.g., CAMK2A, Thr286, Ser314; CAMK2B, Thr287), some phosphorylated peptides were reduced to a much smaller degree (e.g., CAMK2A, Ser279, Ser333/334; CAMK2B, Ser276/Thr277). Interestingly, levels of the unique CAMK2G-or CAMK2D-derived phosphopeptides were not significantly affected (**Figure 2D**).

**Figure 2:**
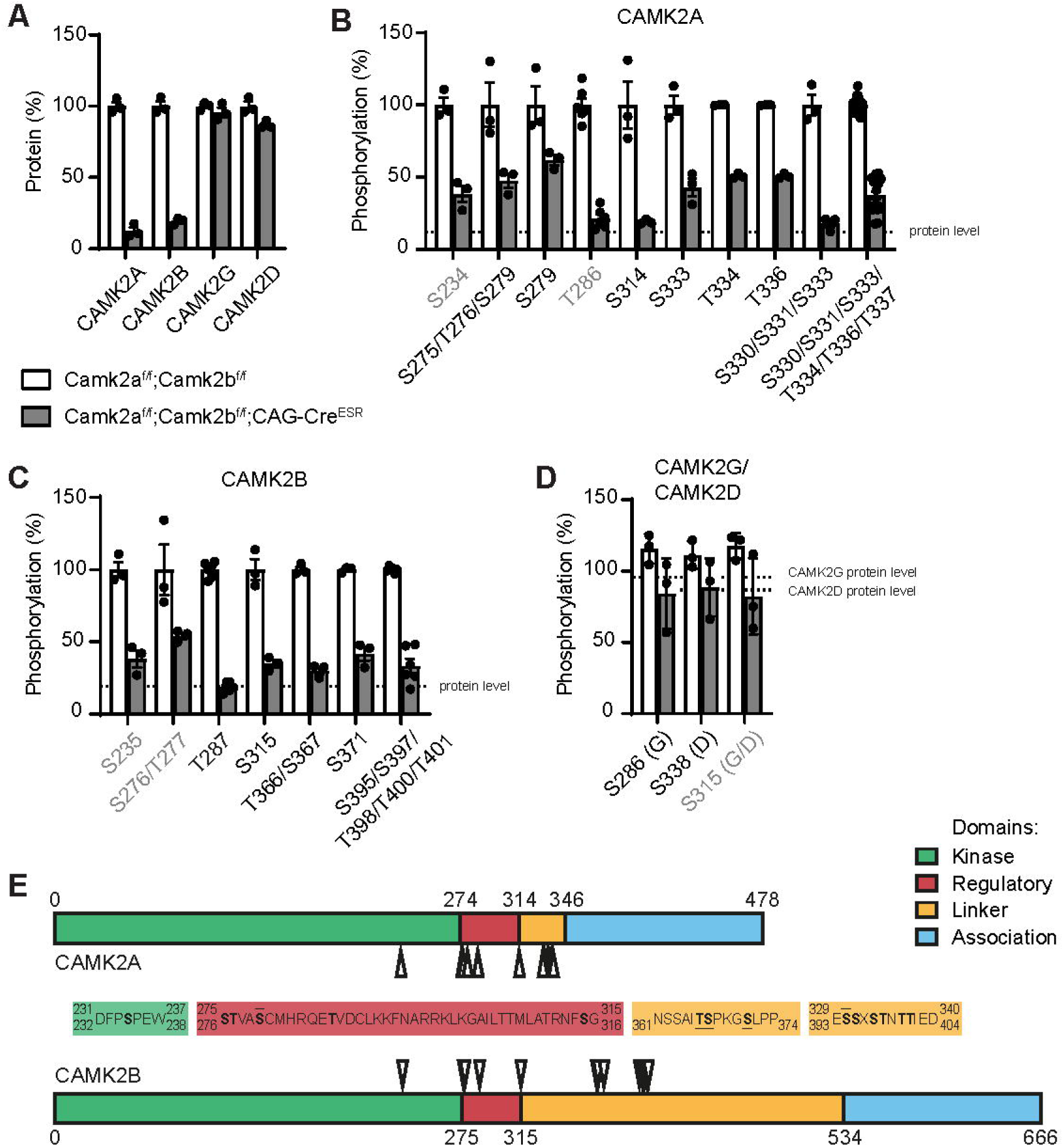
Phosphorylation sites on CAMK2. **(A)** Proteomics analysis showed successful deletion of CAMK2A and CAMK2B but not the other isozymes CAMK2G or CAMK2D in *Camk2a^f/f^;Camk2b^f/f^;Cag-Cre^ESR^* mice. Data is derived from previously obtained dataset (Kool et al., 2019). **(B)** Phosphopeptides identified on CAMK2A are downregulated in *Camk2a^f/f^;Camk2b^f/f^;Cag-Cre^ESR^*mice. Phosphosites in grey are not unique and can belong to more than one CAMK2. The peptide of S234 also corresponds to CAMK2B, CAMK2G and CAMK2D. The peptide of T286 to CAMK2D. The dashed line indicates the CAMK2A protein level in *Camk2a^f/f^;Camk2b^f/f^;Cag-Cre^ESR^* mice at 12%. **(C)** The phosphopeptides identified on CAMK2B are also downregulated in *Camk2a^f/f^;Camk2b^f/f^;Cag-Cre^ESR^*mice. Phosphosites in grey are not unique and can belong to more than one CAMK2. The peptide of S235 also corresponds to CAMK2A, CAMK2G and CAMK2D. The peptide of S276/T277 to CAMK2G and CAMK2D. The dashed line indicates the CAMK2B protein level at 19%. **(D)** Unique phosphosites of CAMK2G (G) and CAMK2D (D) are not downregulated in *Camk2a^f/f^;Camk2b^f/f^;Cag-Cre^ESR^*mice. The dashed lines indicate the CAMK2G and CAMK2D protein levels at 96% and 87%, respectively. **(E)** CAMK2A and CAMK2B linear protein with functional domains on scale. Phosphosites are indicated by arrowheads and are bold in the amino acid sequence. Lines above or below the sequence indicate phosphosites that were only identified in CAMK2A or CAMK2B, respectively.

### CAMK2 substrate motif

Our extensive list of *in vivo* CAMK2 substrates allowed us to examine amino acid sequences surrounding the phosphorylation sites to identify overrepresented amino acids at each position. We aligned the downregulated phosphosites in our dataset to the 0 position and plotted upstream and downstream amino acids. Arg was the most strongly overrepresented at the -3 position, and Lys was also well represented at this position (**Figure 3A**). Acidic amino acids (Asp and Glu) were strongly represented at the +2 position. We often detected hydrophobic residues at the -5 and +1 position, most often represented by a Leu. When we include -6 due to structural variations at the binding site (Özden et al., 2022), we found the hydrophobic residue at -5 or -6 position in 77% of the cases. Additionally, we rarely observed a charged amino acid at the -2 position. The representations of different residues at positions surrounding these phosphorylation sites is largely consistent with the known consensus motif, which includes Arg at -3, hydrophobic amino acids at -5 and +1, and a non-basic amino acid at -2 (Hudmon & Schulman, 2002; Pearson et al., 1985; White et al., 1998; Yasuda et al., 2022). Thus, our in vivo data help refine the established consensus phosphorylation motif at one important position: the preference for an acidic residue at the +2 position (**Figure 3A-B**). Amino acid sequences surrounding ∼75% of our identified phosphorylation sites match 3 or more out of 5 positions within the consensus sequence. However, some of the identified phosphopeptides were very poor matches to the consensus sequence (**Figure 3C**). For example, the phosphopeptide that was most strongly downregulated in our dataset besides the CAMK2-derived peptides was derived from SHANK3 Ser1510 (QLNKDTRS*LGEEP) which lacks a basic residue at -3, and an acidic residue at +2, and contains only an uncharged residue at -2 position (Thr) and a hydrophobic residue at the +1 position (Leu). However, the peptide contains a hydrophobic residue at the -6 position, instead of -5, and basic residues at the -1 and -4 positions, instead of -2. This poor match to the consensus phosphorylation sequence raised a question about whether CAMK2 directly phosphorylates Ser1510 in SHANK3, or perhaps indirectly affects another kinase responsible for this phosphorylation. Therefore, we tested whether purified CAMK2A directly phosphorylated a GST fusion protein containing residues 1468-1740 from SHANK3 *in vitro*, as well as the effect of mutating this putative phosphorylation site (Ser1511 in the rat protein). We found that CAMK2 robustly phosphorylated this GST fusion protein and that phosphorylation was significantly reduced by the Ser1511Ala mutation (**Figure S2**). These data further support the identification of Ser1510 in mouse SHANK3 (Ser1511 in rat) as a direct CAMK2 phosphorylation site.

**Figure 3:**
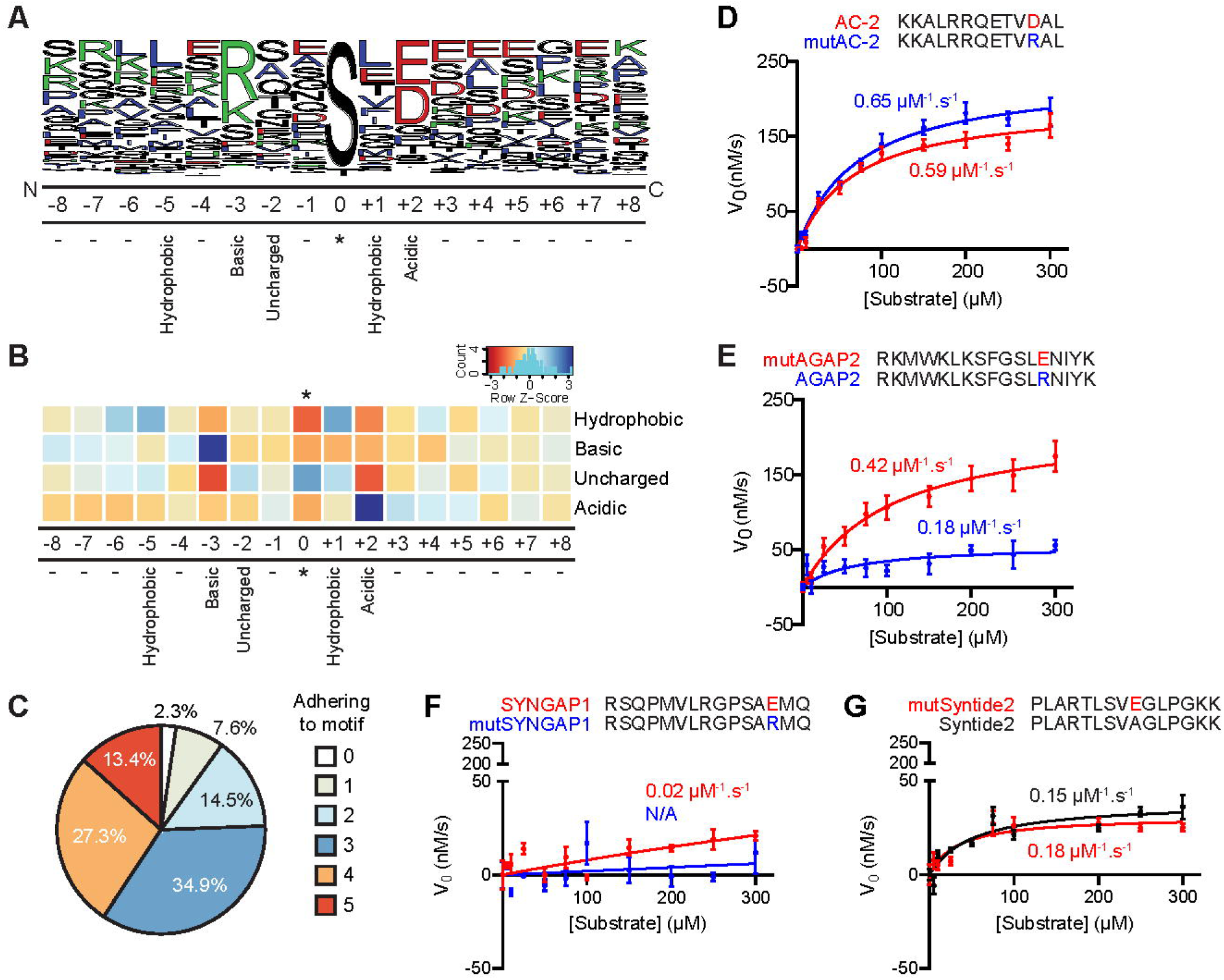
CAMK2 phosphorylation motif. **(A)** Frequency plot of neighboring amino acid (+/-eight) sequence of phosphosites that were set at 0. The motif indicated below is corroborated by literature and our dataset of downregulated phosphosites with confident localization (172 phosphopeptides). Black indicates polar amino acids, green basic, red acidic, and blue hydrophobic. **(B)** Heatmap illustrating the prevalence of the motif on a color scale from red (low) to blue (high), with the amino acid positions below. Note that the uncharged residues showed low prevalence at positions -3 and +2, and high prevalence at positions 0 and +1, but this was due to other rules of the motif. **(C)** Pie chart showing adherence of the phosphopeptides to the motif, i.e. correct residue at -5, -3, -2, +1 and +2 position. **(D-G)** Michaelis-Menten plots for peptide substrates with or without +2 acidic residue: AC-2 (**D**), AGAP2 (**E**), SYNGAP1 (**F**), and Syntide2 (**G**). The phosphorylated residue is shown in bold, and the +2 position is highlighted in red (for acidic residues), blue (for basic residues), or black (for neutral residues).

The acidic residue at the +2 position has been described before (Ando et al., 1991; Hornbeck et al., 2015), but it was not part of the prior CAMK2 consensus motif and its significance has never been tested. To test the effect of the +2 position on substrate phosphorylation, we first determined that there was no correlation between the presence of an acidic residue at the +2 position and the presence of a basic residue at the -3 or -4 position in our dataset of potential CAMK2 substrates (Χ^2^ (1, n = 172) = 0.18, p = 0.669). Next, we performed *in vitro* kinase assays using peptides derived from established CaMKII substrates that natively harbor an acidic residue at +2 position (AC-2 and SYNGAP1) and from those that do not (AGAP2 and Syntide2). For both groups of peptides, we compared the ability of the CAMK2A kinase domain to phosphorylate wild type (WT) peptide versus mutated versions in which the +2 acidic residue was substituted to an Arg (mutAC-2 and mutSYNGAP1) or the +2 was substituted to an acidic residue (mutAGAP2 and mutSyntide2) (**Figure 3D-G**). Out of the four substrates we tested, only AGAP2 phosphorylation was substantially affected by the substitution at the + 2 position (**Figure 3E**). Although we observed a higher K_m_ value for mutAGAP2 (contains a +2 acidic residue), V_max_ and k_cat_/K_m_ values increased ∼4 fold and ∼2.3 fold, respectively (**Table 1**). For the SYNGAP1 peptide, we observed low phosphorylation of the WT sequence, and mutating the +2 position into an Arg completely blocked substrate phosphorylation (**Figure 3F**). For AC-2 and Syntide2, we observed minimal changes in the enzyme kinetics (**Figure 3D, G**).

**Table 1:** Enzyme kinetic parameters for +2 position

| *Substrate* | +2 *aa* | $K_m$<br>($\mu M$) | $V_{max}$<br>($nM/s$) | $k_{cat}$<br>($s^{-1}$) | $k_{cat}/K_m$<br>($\mu M^{-1}.s^{-1}$) |
| --- | --- | --- | --- | --- | --- |
| AC-2 | D (wt) | 65.7 | 194.6 | 38.9 | 0.59 |
|  | R | 71.4 | 232.2 | 46.4 | 0.65 |
| AGAP2 | E | 107.0 | 222.5 | 44.5 | 0.42 |
|  | R (wt) | 61.9 | 56.0 | 11.2 | 0.18 |
| SYNGAP1 | E (wt) | 1446 | 132.2 | 26.4 | 0.02 |
|  | R | N/A | N/A | N/A | N/A |
| Syntide2 | E | 33.5 | 31.0 | 6.2 | 0.18 |
|  | A (wt) | 51.2 | 38.7 | 7.7 | 0.15 |

### Functions of CAMK2 substrates

To get more insight to CaMK2 signaling, we performed a GO analysis on the downregulated phosphoproteins (**Figure 4A**). Significantly enriched terms, such as postsynaptic specialization, glutamatergic synapse and postsynaptic organization, underlined the importance of CAMK2 for glutamatergic signaling in the postsynaptic density. Next, we separated the proteins into functional categories (**Figure 4B**). Many proteins served as scaffolds in the presynapse or postsynapse, supported the cytoskeleton or were regulators of GTPases. We identified phosphopeptides of seven kinases and two phosphatases, whose activity could contribute to alterations in phosphorylation in our database. Interestingly, 26 of the proteins that we identified as putative CAMK2 substrates are associated with neurodevelopmental disorders (NDD), suggesting convergence of NDD pathways (**Figure 4B**). Among the voltage-gated channels and receptors, we only found four potential substrates (GRIN2A, GRIN2B, CACNA1B and KCNQ5). The online resource for post-translational modifications PhosphoSitePlus (Hornbeck et al., 2015) lists many of the phosphosites that we described in our dataset. However, many of them were not yet known to be regulated by CAMK2A or CAMK2B. A comparison to known substrates of CAMK2A and CAMK2B reveals that our dataset provides proof for 113 novel potential substrates of CAMK2 (**Figure 4C**). Our dataset and PhosphoSitePlus share 18 phosphoproteins, yet we find novel phosphopeptides on 11 of these known substrates (LRRC7, MYH9, GRIN2B, RIMS1, SPTBN4, SYNGAP1, DLGAP and SHANK proteins). The PhosphoSitePlus CAMK2 substrates were found with different techniques and predominantly in human, mouse or rat models (**Figure 4D-E**). Therefore, our dataset is a meaningful mouse *in vivo* resource that unveils many additional CAMK2-dependent signaling pathways.

**Figure 4:**
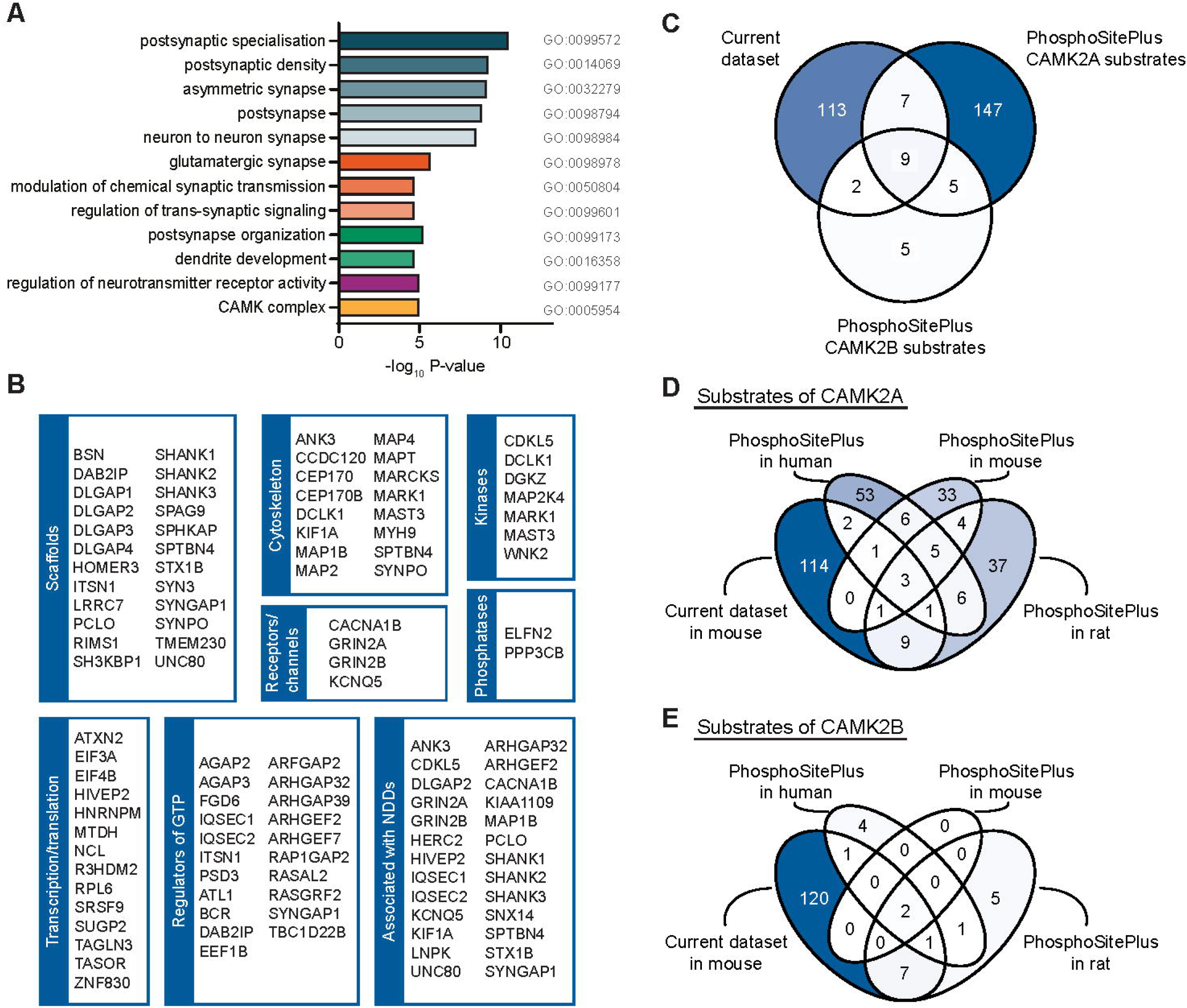
Potential substrates of CAMK2. **(A)** Significantly enriched GO terms after analysis of downregulated phosphoproteins over all identified phosphoproteins, clustered by color. **(B)** Selection of potential substrates of CAMK2 categorized by protein function and association with neurodevelopmental disorders (NDDs). **(C)** Comparison of phosphoproteins identified in our dataset and known substrates of CAMK2A and CAMK2B, derived from the PhosphoSitePlus^®^ database. **(D)** Comparison to the PhosphoSitePlus^®^ substrates of CAMK2A, grouped by species. **(E)** Comparison to the PhosphoSitePlus^®^ substrates of CAMK2B, grouped by species.

## Discussion

CAMK2A and CAMK2B are among the most abundant kinases in the brain and share a high homology. Deleting either gene in mice results in cognitive impairments yet deleting both genes simultaneously is lethal, indicating at least a partial functional redundancy between the two isozymes and suggesting that there still are functions of CAMK2 to be uncovered. To gain insight in these novel functions of CAMK2, we used cortical tissue of inducible *Camk2a^f/f^;Camk2b^f/f^;CAG-Cre^ESR^* mice for phosphoproteomics analysis. With our dataset, we confirm already known substrates *in vivo* and we provide a list of many novel potential substrates of CAMK2. Furthermore, our dataset confirms the important role of CAMK2 in synaptic plasticity and other neuronal signaling pathways.

The lethality of *Camk2a^-/-^Camk2b^-/-^* mice might be explained by changes in protein or phosphorylation levels. Few proteins were up or downregulated in our proteomic analyses in *Camk2a^f/f^;Camk2b^f/f^;CAG-Cre^ESR^* mice (Kool et al., 2019). Only PPLM1L has been associated with increased mortality, but this could be explained by the inability of *Ppm1l*-deficient moms to produce milk for their pups (Kusano et al., 2016; Ren et al., 2014). However, we identified multiple downregulated phosphoproteins that result in lethality when knocked out, including SYNGAP1, ANK3, UNC80, KIF1A, MAP1B, MARCKS, GRIN2B, KCNQ5, ZNF830, ARHGAP39, ARHGEF7, KIAA1109 and HERC2 (Blake et al., 2021). A loss-of-function effect due to lack of phosphorylation by CAMK2 on any of these substrates could potentially explain why *Camk2a^-/-^Camk2b^-/-^* and *Camk2a^f/f^;Camk2b^f/f^;CAG-Cre^ESR^* mice die. It remains an open question whether one of these substrates alone leads to lethality or whether it is a cumulative effect of multiple substrates.

Previous studies have shown that in wild type mice or rats, CAMK2A and CAMK2B are partially autophosphorylated at Thr286 and Thr287, respectively, under steady state (basal) conditions, consistent with the measurement of significant autonomous CAMK2 activity in brain extracts (Lundby et al., 2012). We describe here for the first time an extensive list of proteins with significant decreases in levels of phosphorylation at multiple sites when CAMK2A and CaMK2B are knocked out, representing potential direct CAMK2 substrates at basal state *in vivo*. Approximately 14.5% of previously identified CAMKAPs (Baucum et al., 2015), were identified as basal substrates of CAMK2. The list of phosphopeptides that we detected include known substrates such as scaffolding proteins, GTPase regulatory proteins and NMDA receptor subunits (Dosemeci & Jaffe, 2010; Funahashi et al., 2024; Mizutani et al., 2008; Mota Vieira et al., 2020) but also many proteins whose phosphorylation was not yet known to (also) depend on CAMK2. It is noteworthy that we do not find a big overlap (only ∼15%) between the phosphoproteins in our dataset and CAMK2 substrates listed on the PhosphoSitePlus database. PhosphoSitePlus is a resource developed by Cell Signaling Technology that integrates data from published articles with unpublished data generated by Cell Signaling Technology. The lack of overlap with our findings could partially be explained by the fact that the majority of CAMK2 substrates in PhosphoSitePlus were identified *in vitro.* In this regard, our study of cortical tissue shows overlap with approximately one third of substrates recently identified in striatal slices Click or tap here to enter text.following stimulation of NMDA receptors or voltage-gated calcium channels (Funahashi et al., 2024). In contrast, we identified *in vivo* substrates without stimulating cortical tissue based on reduced phosphorylation in double *Camk2a^f/f^;Camk2b^f/f^;CAG-Cre^ESR^* knockout mice. This implies that we identify substrates that are not necessarily phosphorylated only upon stimulated conditions, but already are phosphorylated in basal state. We hypothesize that these substrates might therefore be required for homeostatic purposes, which might be linked to maintenance of an intrinsic memory. Adult deletion of CAMK2A and CAMK2B simultaneously disrupts this homeostatic maintenance, potentially causing the premature death of the mice. Further research is required to obtain more evidence for this hypothesis.

The strongest downregulated phosphopeptide in *Camk2a^f/f^;Camk2b^f/f^;CAG-Cre^ESR^*was on SHANK3, whose activity-dependent interaction with CAMK2A has been a topic of recent interest (Cai et al., 2021; Perfitt, Wang, et al., 2020; Tao-Cheng, 2020). The specific residue (Ser1510) was previously reported to depend on CAMK2A *in vitro* (Dosemeci & Jaffe, 2010), and we confirmed that CAMK2A can directly phosphorylate this site *in vitro*, but the functional effect remains to be studied. Recently, another phosphorylated residue on SHANK3 (Ser685) was shown to be phosphorylated by CAMK2 *in vitro* (Perfitt, Stauffer, et al., 2020). We did not find SHANK3 at Ser685 in our list of phosphopeptides, which might be due to the fact that this phosphosite is not present in basal state. In addition, a PubMed search (November 2024) failed to detect any prior reports of CAMK2 phosphorylation of >65% of the phosphoproteins that were detected in our study, representing previously unrecognized putative substrates of CAMK2. Examples are the phosphatase ELFN2, potassium channel KCNQ5, and many GTP regulating proteins such as AGAP and DAB2IP. This list provides avenues for future research to uncover the impact of CAMK2 on the pathways these proteins are involved in.

CAMK2 plays an important role in the synapse, which is highlighted by the GO pathway analysis of the substrates. Many proteins that are important in the synapse are associated with neurodevelopmental disorders, or synaptopathies, such as scaffolds SHANK and DLGAP, voltage gated ion channels KCNQ5 and CACNA1B, and GTP regulators IQSEC and SYNGAP1. CAMK2A and CAMK2B have also been associated with neurodevelopmental disorders (Akita et al., 2018; Chia et al., 2018; Iossifov et al., 2014; Küry et al., 2017; Rizzi et al., 2020; Stephenson et al., 2017). Our dataset can hopefully enhance the understanding of the functional pathways of these synaptic proteins, ultimately improving potential treatment for patients with synaptopathies. However, it is important to mention that despite the preponderance of synaptic proteins detected in this study, we failed to detect some well-known synaptic CAMK2 substrates, such as GluA1, perhaps because of their relatively low abundance or inefficient enrichment of these phosphopeptides using our methodology. Thus, our findings do not exclude the possibility that additional CAMK2 substrates remain to be detected.

Phosphorylation residues that we found on CAMK2A and CAMK2B include the autophosphorylation residues Thr286 or Thr287, respectively, but not the next best studied autophosphorylation residues Thr305/Thr306 or Thr306/Thr307, respectively. The phosphorylation of these last two residues is strongly enhanced following CAMK2 activation, whereas the tissue was harvested under baseline conditions for our studies. However, prior studies using phospho-specific antibodies detected phosphorylation of CAMK2A at Thr305/Thr306 in the hippocampus (Elgersma et al., 2002), and a prior phosphoproteomic study detected CAMK2A Thr306 phosphorylation in samples isolated from mouse forebrain (Baucum et al., 2015). One potential explanation for this discrepancy is that levels of CAMK2A Thr306 phosphorylation may be lower in cortex that in other part of the forebrain. Interestingly, the levels of the various phosphorylated CAMK2A and CAMK2B peptides that we detected were not consistently decreased in parallel with the 80-90% decrease in levels of total CAMK2A and CAMK2B protein. Although levels of CAMK2A phosphorylation at Thr286 and Ser314 also decreased by 80-80%, the phosphorylation of Ser275 and Thr333 in CAMK2A was reduced by only ∼50%. Although the function(s) of CAMK2A phosphorylation at these sites is unknown, it can be hypothesized that the relative levels of phosphorylation at these sites is increased in the face of the loss of total CAMK2 protein in double knockout mice in order to preserve an unknown essential CAMK2 function. Interestingly, previous research has shown that phosphorylation at Ser275, Thr276 and Ser279, shows less than 10% increase upon calcium stimulation *in vitro* (Baucum et al., 2015) and these phosphorylation sites were not identified in the recent *ex vivo* study in striatum where they compared stimulated with non-stimulated tissue (Funahashi et al., 2024). This further supports our hypothesis that phosphorylation at some substrates is not stimulation dependent. One caveat to consider is that with the low CAMK2 levels in the *Camk2a^f/f^;Camk2b^f/f^;CAG-Cre^ESR^*samples, the error margin is increased.

Our data provide further validation of the CAMK2 consensus motif to consist of -5 hydrophobic, -3 basic, +1 hydrophobic and +2 acidic amino acid, aligning the phosphorylated residue at position 0. We show that the position at +2 has a conditional effect that is substrate specific. The +2 acidic residue can substantially increase catalytic efficiency for some substrates whereas in others it does not have a notable effect, at least in the context of synthetic peptide substrates. It seems clear that substrate specificity and catalytic efficiency are determined by a combination of interacting residues around the phosphorylation site, where if some sites are not satisfied, then having an acidic residue at +2 will significantly impact catalysis. But if all other positions are satisfied, it is not necessary to satisfy the +2 acidic residue for efficient binding and phosphorylation. We show Leu as the main residue at the +1 position, which most likely regulates the Ser/Thr specificity of CAMK2 (Johnson et al., 2023). Very recently a binding consensus sequence was identified for CAMKAPs to interact with CAMK2 (Özden et al., 2022). The binding consensus sequence overlaps with the phosphorylation consensus motif described here, with a hydrophobic residue situated at the - 5 and +1 position, and a basic residue at -3. The same group described a Gln at -2, which we also found as the third most frequent residue, and a hydrophobic amino acid at -8, which was less present in our motif. However, it seems that the motif for binding of or phosphorylation by CAMK2 is highly similar and that phosphorylation or stable binding to this region is potentially determined by whether or not there is a serine or threonine present at the 0 position. Also, the motif of CAMK2D, tested in kidney cells, is determined by the basic residue at -3 and the acidic residue at +2 (Park et al., 2023), indicating that these two positions in the consensus motif are shared between different CAMK2 isozymes.

Taken together, our data expand the knowledge of CAMK2 substrates and emphasize its functional importance in synaptic signaling. Further elucidating the pathways CAMK2 is involved in, we provide an extensive list of proteins that lose their phosphorylation site upon deletion of *Camk2a* and *Camk2b*.

## Acknowledgements

This research was supported by the Dutch Research Council (ALW-Veni (863.12.017) and NWO-Vidi (016.Vidi.188.014) to GW).

## Author contributions

PR performed the experiments and analyses; JD performed the mass spectrometry; OK performed enzyme kinetics experiments; TP performed *in vitro* Shank3 phosphorylation studies, GvW, PR, MS, TP and RC designed the experiments; RC, MS and YE gave critical input; PR and GvW drafted the manuscript, and all authors contributed to the final version of the manuscript.

## Data availability

The mass spectrometry raw files were uploaded to PRIDE/EBI and once a PXD identifier has been allocated it will be added to the manuscript text.

---

**Supplemental figure 1.**
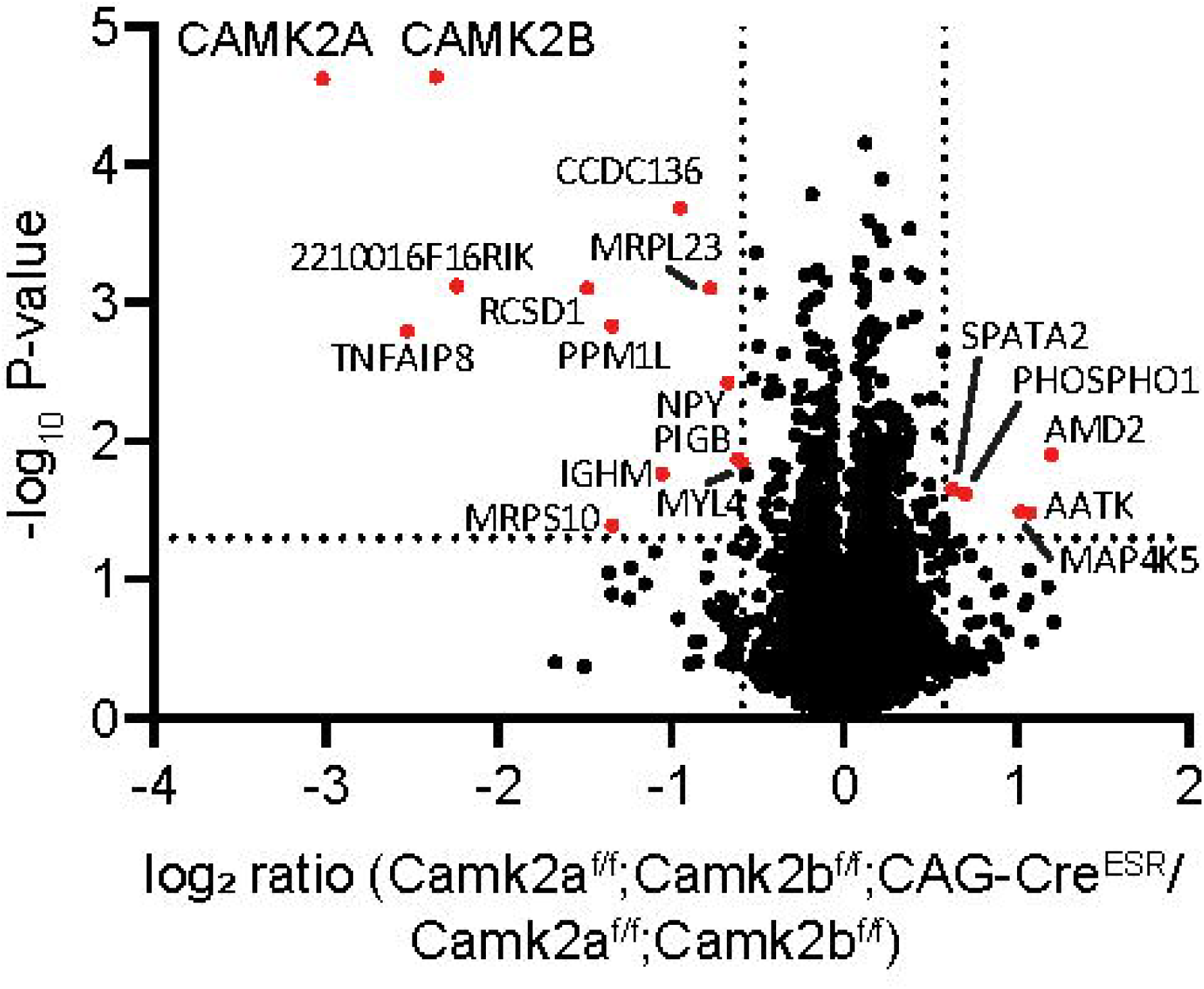
Volcano plot of the proteomics dataset, showing minimal changes on protein level when *Camk2a* and *Camk2b* are deleted. Red dots indicate proteins that are signficantly up-or downregulated >1.5 fold in *Camk2a’^11^;Camk2bvr;­* CAG-CreEsR compared to *Camk2a’’.·Camk2b’^1^’* mice. Data is derived from previously obtained dataset (Kool et al.. 2019).

**Supplemental figure 2.**
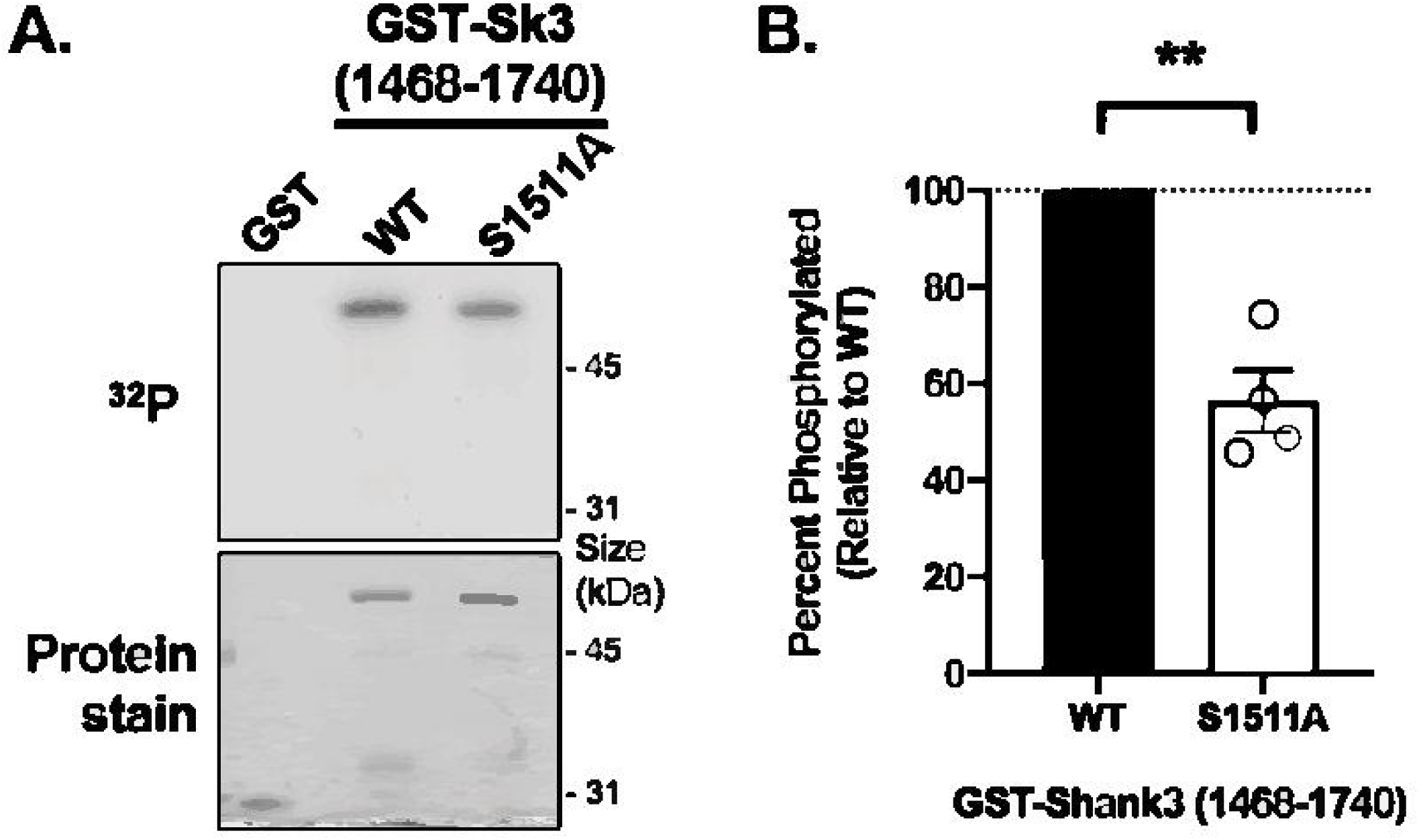
A) Representative Coomassie Blue stained gel and autoradiograph from 1 of 4 replicates. B) Bar graph with super-imposed scatter plot summarizing quantified data from 4 independent replicates. The S1511Amutation (rat) significantly reduced phosphorylation of GST-Sk3(1468-1740) by 43.7±6% compared to WT (** p = 0.006, one-sample Student’s I-test with equal variance compared to a theoretical value of 100).

